# Fast fluorescence lifetime imaging reveals the aggregation processes of α-synuclein and polyglutamine in aging *Caenorhabditis elegans*

**DOI:** 10.1101/414714

**Authors:** Romain F. Laine, Tessa Sinnige, Kai Yu Ma, Amanda J. Haack, Chetan Poudel, Peter Gaida, Nathan Curry, Michele Perni, Ellen A.A. Nollen, Christopher M. Dobson, Michele Vendruscolo, Gabriele S. Kaminski Schierle, Clemens F. Kaminski

## Abstract

The nematode worm *Caenorhabditis elegans* has emerged as an important model organism to study the molecular mechanisms of protein misfolding diseases associated with amyloid formation because of its small size, ease of genetic manipulation and optical transparency. Obtaining a reliable and quantitative read-out of protein aggregation in this system, however, remains a challenge. To address this problem, we here present a fast time-gated fluorescence lifetime imaging (TG-FLIM) method and show that it provides functional insights into the process of protein aggregation in living animals by enabling the rapid characterisation of different types of aggregates. More specifically, in longitudinal studies of *C. elegans* models of Parkinson’s and Huntington’s diseases, we observed marked differences in the aggregation kinetics and the nature of the protein inclusions formed by α-synuclein and polyglutamine. In particular, we found that α-synuclein inclusions do not display amyloid-like features until late in the life of the worms, whereas polyglutamine forms amyloid characteristics rapidly in early adulthood. Furthermore, we show that the TG-FLIM method is capable of imaging live and non-anaesthetised worms moving in specially designed agarose micro-chambers. Taken together, our results show that the TG-FLIM method enables high-throughput functional imaging of living *C. elegans* that can be used to study *in vivo* mechanisms of aggregation and that has the potential to aid the search for therapeutic modifiers of protein aggregation and toxicity.

---

A variety of human diseases, including neurodegenerative disorders such as Parkinson’s and Alzheimer’s diseases, are characterised by the misfolding of protein species and their subsequent aggregation into amyloid fibrils_1,2_. The nematode *Caenorhabditis elegans* is a particularly useful model organism through which to study these diseases^3–8^ and to screen for small molecule inhibitors of the protein aggregation process_9,10_. *C. elegans* has a simple body plan of 959 somatic cells, its genetics are well-characterised, and at least 40% of its genes have known human homologs_11_. Furthermore, it has a relatively short lifespan of only 2-3 weeks and is optically transparent, making it a highly suitable system for longitudinal imaging studies of protein aggregation. Despite these advantages, obtaining quantitative read-outs for amyloidogenic protein aggregation *in vivo* remains challenging_12_. Fluorescence intensity measurements are prone to artefacts, and classifying aggregates by counting protein inclusions relies on arbitrary choices for intensity and size cut-offs. Furthermore, reliable discrimination between amyloid-like and amorphous aggregates is generally not possible. The use of thioflavin T, a dye that becomes fluorescent on intercalation into the cross β-structure of amyloid aggregates, and that is commonly used for the study of protein aggregation *in vitro*, is not compatible with live worm imaging because it affects protein homeostasis in the nematodes_13_.

To address some of these issues, we have previously established a readout for the state of protein aggregation based on fluorescence lifetime imaging microscopy (FLIM) of a fluorophore covalently linked to the amyloidogenic protein of interest_14_. FLIM not only informs on the location but also on the molecular environment of the fluorescent probes, providing fully quantitative read-outs^15–17^. We have shown that a reduction in lifetime from the reporter fluorophore correlates with the degree of aggregation of the protein to which it is attached, and that this provides a quantitative measure of the degree of protein aggregation *in vitro*, in live cells and in C*. elegans*_14_. The decrease in lifetime is thought to be associated with fluorescence energy transfer to intrinsic energy states associated with the amyloid fibrils_14_. Conjugated organic fluorophores^18–20^ and intrinsic protein fluorescence_21_ have also been used successfully as FLIM sensors for protein aggregation, as has the amyloid-binding dye heptamer-formyl thiophene acetic acid (hFTAA)_22_. Conventional FLIM measurements are slow, however, as they are based on time-correlated single photon counting (TCSPC)_23,24_ which involves acquisition times on the order of 2 min for a single field of view. This method therefore requires the use of anaesthetised or fixed animals, greatly limiting the throughput of the technique and preventing studies of freely moving, live animals.

In this work, we specifically set out to address these problems and to establish a method that improves throughput and physiological relevance and permits studies of moving animals. The method makes use of time-gated FLIM (TG-FLIM)_25_, a fast and quantitative imaging modality which provides unprecedented throughput for FLIM measurements of non-paralysed animals. Unlike TCSPC, which is usually performed in conjunction with laser scanning confocal microscopy (LSCM), TG-FLIM is a wide-field technique and thus heavily parallelises the FLIM measurements. The fluorescence decay is measured by collecting the fluorescence signal with nanosecond-wide temporal gates, which are temporally shifted to enable the fluorescence decay to be sampled. Gating and time shifting are achieved with a high-rate imager (HRI), and lifetime data are thus obtained in each pixel from a sequential set of images gated at different time delays (**Figure 1a**). TG-FLIM has been successfully used for high-throughput imaging of protein-protein interactions and biosensors in cells_26,27_, but not so far in the context of protein aggregation, or in worm models of disease. Here we describe a novel approach for aggregation studies in *C. elegans* models of protein misfolding diseases based on TG-FLIM. We show that we can monitor intracellular aggregation in the worms over their entire lifespan using TG-FLIM with excellent repeatability and precision in the measured lifetimes. Our approach reveals differences in the kinetics of protein aggregation and the type of species appearing in the worms for two disease models. Furthermore, we provide details of the quantification of TG-FLIM data from live, moving animals and introduce motion correction algorithms for the TG-FLIM data analysis.

## Results and discussion

### Time-gated FLIM provides a robust lifetime readout from *C. elegans*

A schematic of the TG-FLIM microscope setup that we have developed is shown in **Figure 1a**, details of which are found in the Methods section and in **Supplementary Figure 1**. To assess the precision and repeatability of the method, we measured the fluorescence lifetimes of yellow fluorescent protein (YFP) when expressed in *C. elegans*. **Figure 1c** shows the head region of worms expressing YFP in body wall muscle cells. For the purpose of these experiments, the worms were anaesthetised. The average standard deviation across pixels within a single field of view was 17 ps (as shown by the error bars on **Figure 1d)**. The variability between individual worms was estimated by comparing the mean lifetimes obtained from the worms and found to be 16 ps (standard deviation). The system exhibited excellent intra-frame and inter-frame repeatability of the value of the fluorescence lifetimes measured in this way. The results presented in **Figure 1c,d** were obtained by acquiring 61 equally-spaced gates with exposure times of 65 ms each. The gate width was set to 1 ns and the time gated images were acquired every 250 ps, leading to a total acquisition time of 4.2 s per field of view. This corresponds to a ~30-fold improvement in speed compared to typical TCSPC measurements of similar quality_14_, thus allowing for a much higher throughput. The results, therefore, show that TG-FLIM is a robust and quantitative readout for biosensors in *C. elegans*, featuring good spatial and temporal resolution compatible with high-throughput or dynamic imaging.

**Figure 1.**
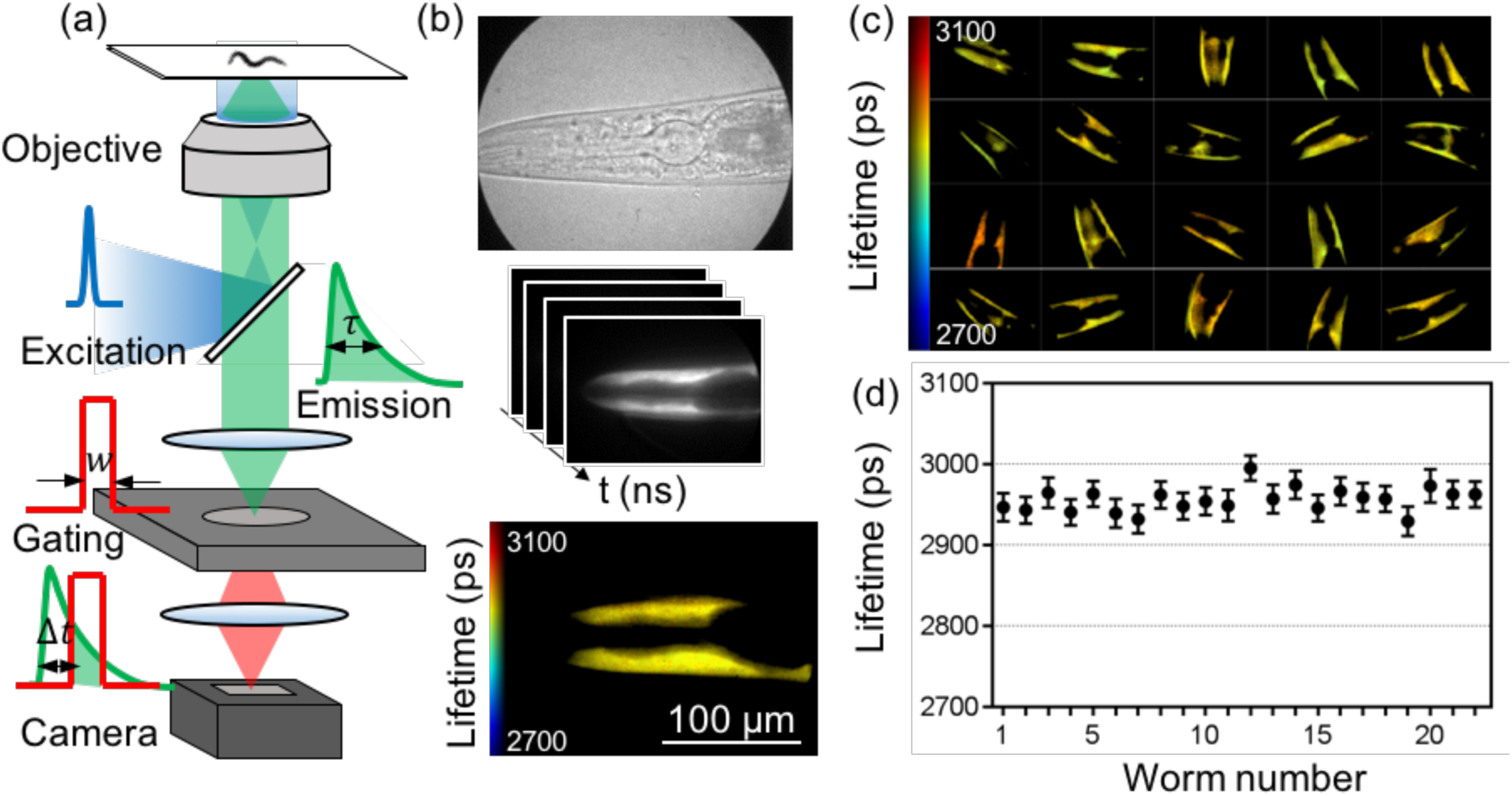
TG-FLIM imaging of live *C. elegans* expressing YFP in the body wall muscle cells. **(a)** Schematic of the FLIM setup, showing the excitation pulse (blue curve), the fluorescence emission decay (green curve) and the gated detection (red curve) performed by the high-rate imager (HRI). The fluorescence lifetime (τ) is measured by gating the fluorescence decay with a gate width (*w*) and a set of gate positions (Δ*t*). **(b)** Bright-field image acquired with the HRI camera (top), schematic FLIM stack consisting of images recorded at different time gates (middle), and the reconstructed fluorescence lifetime map obtained through such measurements from a single worm head region (bottom). **(c)** Fluorescence lifetime maps of 20 individual worms. Each field of view represents a 225 μm x 167 μm region in the sample plane. **(d)** Average lifetimes obtained from individual worms demonstrating the precision (error bars are standard deviation across pixels within the field of view) and the variability of lifetimes between individual worms in one experiment.

### Longitudinal FLIM studies of Parkinson’s and Huntington’s disease models

We then explored if the method is capable of detecting age-associated protein aggregation in *C. elegans* models of protein misfolding diseases. To this end, we carried out a longitudinal study of the fluorescence lifetimes and the distribution of the aggregates that form in *C. elegans* models for Parkinson’s and Huntington’s diseases, expressing YFP-tagged variants of α-syn_5_ and polyglutamine (40 glutamine residues, Q40)_4_, respectively, in the body wall muscle cells. We focused on imaging the head region of the worms and obtained sufficient resolution to be able to identify individual inclusions. Data were recorded across the whole lifespan of a population of nematodes, using the strain expressing only YFP as a control. We imaged a pool of ca. 20 worms for each strain (ca. 60 worms in total) on days 0, 3, 6, 10, 12 and 14 of adulthood for two independent biological replicates. The fluorescence lifetimes for each population and representative TG-FLIM images are shown in **Figure 2**.

**Figure 2.**
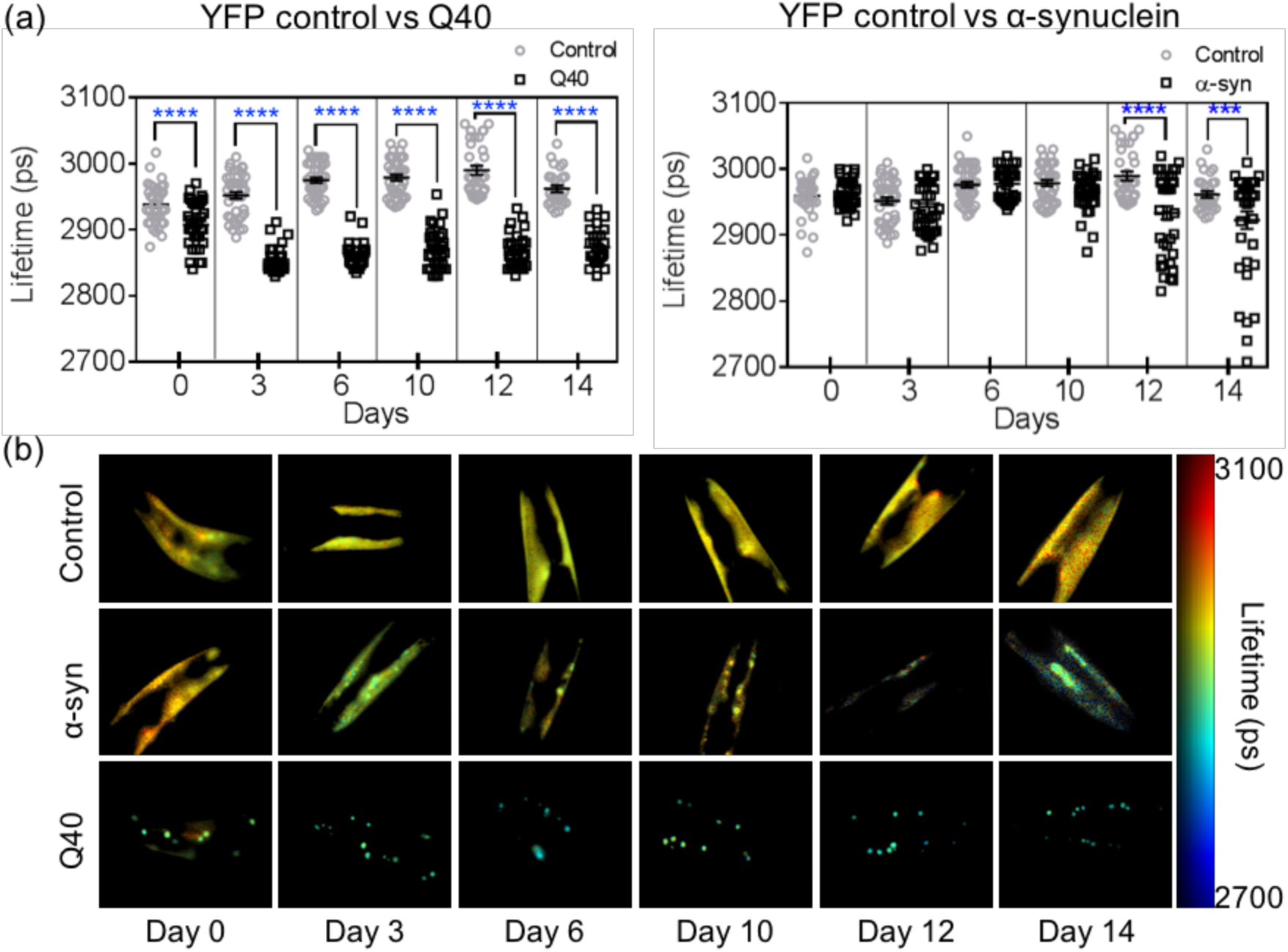
TG-FLIM imaging of protein aggregation in Q40 and α-syn strains across the lifespan of the animals. **(a**) Fluorescence lifetimes at day 0, 3, 6, 10, 12, and 14 of adulthood for Q40 and α-syn compared to YFP controls. Squares and circles represent the average lifetime of the fluorescence from the head region of each imaged worm. Statistical analysis was performed using a one-way ANOVA, **** p<0.0001. The data shown are pooled from two biological replicates totalling ca. 40 worms per strain and time point. **(b)** Representative worm FLIM images for each day and each strain.

We observed that the average fluorescence lifetime of Q40 dropped early in the life of the worms (between day 0 and day 3) and remained constant throughout the rest of their lifespan (**Figure 2a**). At day 0, the Q40 worms presented a combination of bright inclusions with lower fluorescence lifetimes than the YFP control, indicative of protein molecules being in an amyloid state_14_, as well as diffuse signal with lifetimes similar to that of the YFP control, therefore indicative of a non-amyloid state (**Figure 2b** and **Figure 3a**). The diffuse signal was not observed from day 3 onwards, and we infer that at this stage of the worm life all soluble protein had been incorporated in amyloid-like inclusions in agreement with the significantly lower average lifetimes of ca. 2850 ps compared to the YFP control of ca. 2950 ps (**Figure 2a**). These results are consistent with previous fluorescence recovery after photobleaching (FRAP) studies, where the diffuse Q40 observed in young animals was found to be relatively mobile, compared to the Q40 inclusions which were immobile_4,28_.

In contrast to the Q40 animals, the fluorescence lifetime of α-syn remained similar to that of the YFP control worms until day 10, after which a reduction in lifetime was observed (**Figure 2a**). Detailed analysis revealed that inclusions were present in α-syn worms from day 0 onwards, yet these did not correspond to highly ordered aggregates as judged from the uniform fluorescence lifetimes observed across the animals throughout most of their lifespan (**Figure 2b** and **Figure 3b**). Only late in the life of the animal, after day 10, did we observe α-syn inclusions with fluorescence lifetimes indicative of well-defined amyloid fibrils (**Figure 2b**). Again, these results are consistent with FRAP data, in which immobile inclusions were observed only from day 11 onwards_5_. The precise molecular nature of the earlier inclusions is unclear at present, but we speculate that they could either be disordered oligomeric species as observed in the early stages of α-syn aggregation in previous studies_29,30_, accumulations of monomeric α-syn with lipids or other cellular components, or perhaps droplets with liquid-like properties which have recently attracted attention as possible precursors of amyloidogenic protein _31_ aggregation.

**Figure 3.**
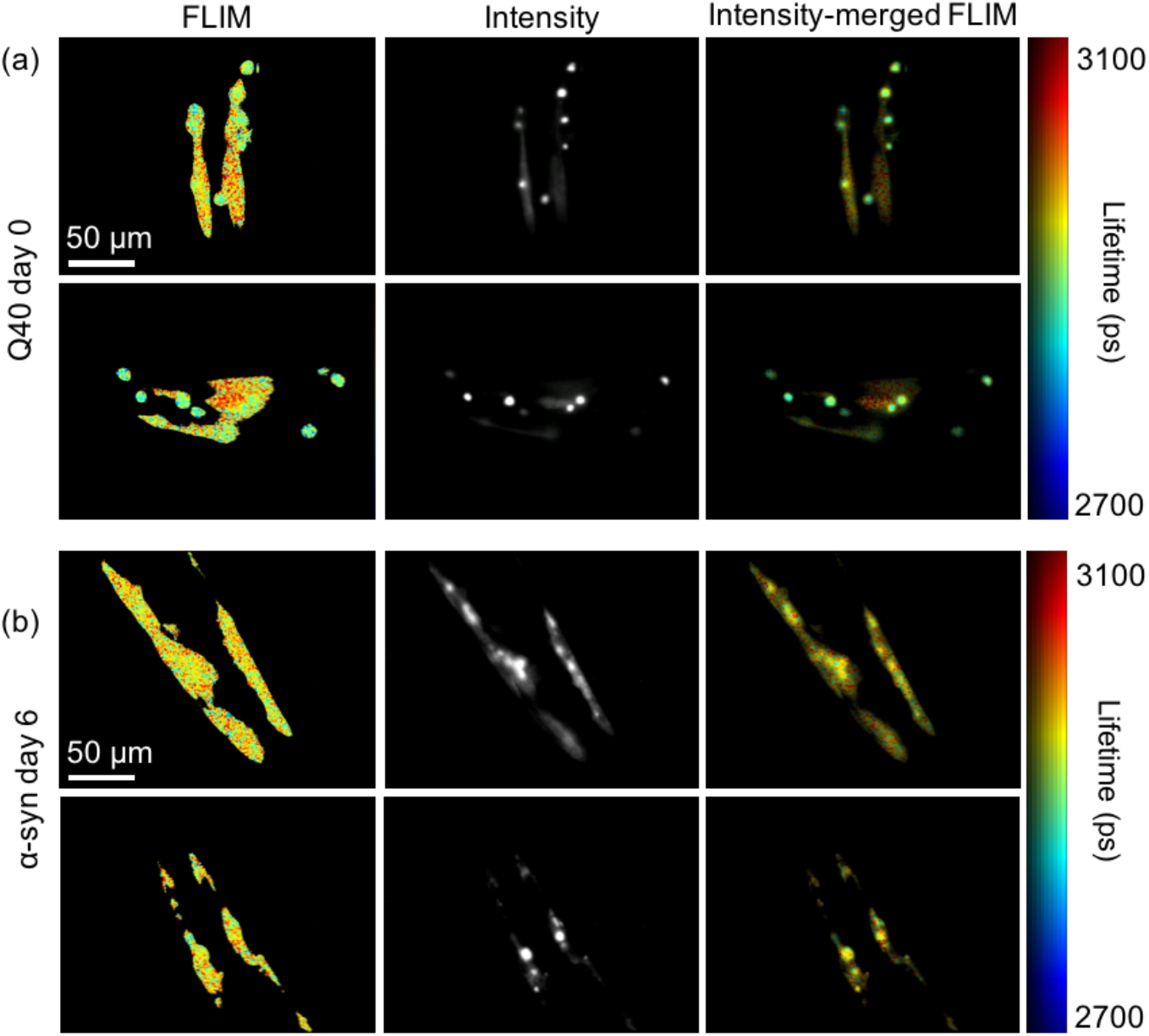
Distribution of Q40 at day 0 (**a**) and α-syn at day 6 (**b**) highlighting the different aggregation states. Shown are the false-colour FLIM maps (left), the signal intensity (middle) and the intensity-merged FLIM maps (right). **(a)** Representative images of Q40 worms at day 0. The FLIM and intensity maps show that the protein is distributed between diffuse signal and foci, the latter having a shorter lifetime. **(b)** Representative images of α-syn worms at day 6 presenting bright inclusions with comparable lifetime as the diffuse signal.

Thus, this longitudinal TG-FLIM study reveals clear differences in the kinetics of the aggregation process for strains expressing Q40 and α-syn, and informs on the nature of the protein inclusions *in vivo* as the animals age.

### TG-FLIM imaging of live *C. elegans* crawling in agarose micro-chambers

Imaging studies of *C. elegans* are typically performed on anaesthetised animals in order to circumvent motion artefacts at the examined length-and time-scales. Although the animals remain alive during the experiment, the use of anaesthetics is not compatible with the observation of behavior, nor does it allow an individual animal to be monitored over long periods of time. Given the exceptional speed and precision of TG-FLIM, we set out to examine its applicability for imaging live and crawling *C. elegans*. To this end, we designed agarose micro-chambers that can hold individual worms (**Figure 4a**), inspired by previous studies on both *C. elegans* larvae and adults_32,33_. The dimensions of the micro-chambers were chosen so that worms remained within the field of view and the depth of field of the microscope and were able to crawl freely within the micro-chamber as shown in the supplementary video (see **Supplementary Figure 2**).

As a benchmark for this approach, we inserted *C. elegans* expressing YFP and Q40 at day 3 of adulthood in the micro-chambers for TG-FLIM measurements (**Figure 4**). To increase acquisition speed, we used only 7 time gates (gate width of 1 ns with time gates acquired every 2.175 ns) compared to the 61 used for studies described in the preceding sections, thus shortening the total recording time to just 0.5 s per FLIM acquisition and minimising movement artefacts across the TG-FLIM dataset.

Although the worms did not move on the timescale of the individually recorded frames in our TG-FLIM measurements, motion over the entire acquisition sequence (0.5 s) caused significant artefacts during the FLIM reconstruction, as a consequence of the loss of spatial correspondence of image pixels between the different time gates (**Supplementary Figure 3**). In order to correct for the worm movement during the data acquisition, we developed an image registration procedure that re-aligns each dataset before FLIM analysis, similar to that used previously to remove motion artefacts from intravital imaging_34_. The procedure uses a non-rigid transformation to register features in each individual frame compared to the brightest frame of the given TG-FLIM dataset. We quantified the resulting standard deviations on control worms and found a lower lifetime resolution compared to the experiments on anaesthetised animals (higher standard deviation of ca. 60 ps with 7 gates, compared to 17 ps with 61 gates, with a similar level of signal in the maximum time gate). This difference can be explained by a lower sampling rate of the fluorescence decay and therefore a lower number of total photons in the decay, as well as residual errors in registration adding some noise to the lifetime estimation.

Additionally, we implemented a digital “worm stretching” procedure inspired by the work of Christensen *et al*._35_. This approach has two major advantages: first, it allows for the averaging of fluorescence lifetime maps of the same worm if consecutively imaged, improving the signal-to-noise ratio (SNR) of the resulting lifetime map. Second, it allows alignment of images of multiple worms along a similar template (a “stretched” form of the worm), which enables a direct comparison of multiple datasets for the assessment of the distribution of markers and functional read-outs. Being able to visualise data in a tractable way is important especially when a high throughput approach is taken. The digital stretching was performed by first determining the worm outline and drawing the backbone of the worm via skeletonisation, using a similar approach to that used for tracking worms_36_. The backbone was then used to extract the information about the curvature of the worm, which in turn allowed reconstruction of the signal from the digitally stretched worm (see Methods for details).

Representative results of the YFP control and Q40 worms are shown in **Figure 4b**. The lifetime maps of the worms were digitally stretched as described above and 10 consecutive lifetime maps were averaged in order to achieve a comparable image quality to that obtained in our study on the anaesthetised worms. The fluorescence lifetime maps reconstructed here show no visible artefacts despite the motion of the worms during the acquisition. Additionally, we found that the fluorescence lifetimes obtained this way are in good agreement with those obtained in the experiments on immobilised animals (compare **Figure 4b** to **Figure 2b**). These measurements therefore allow functional imaging of aggregate states in entire worms, and enable their behavior to be monitored over time. Furthermore, the visualisation of worms as stretched templates, as in **Figure 4b**, allows a direct comparison of the degree of aggregation (derived from the FLIM measurement) and of the spatial distribution of the protein deposits.

**Figure 4.**
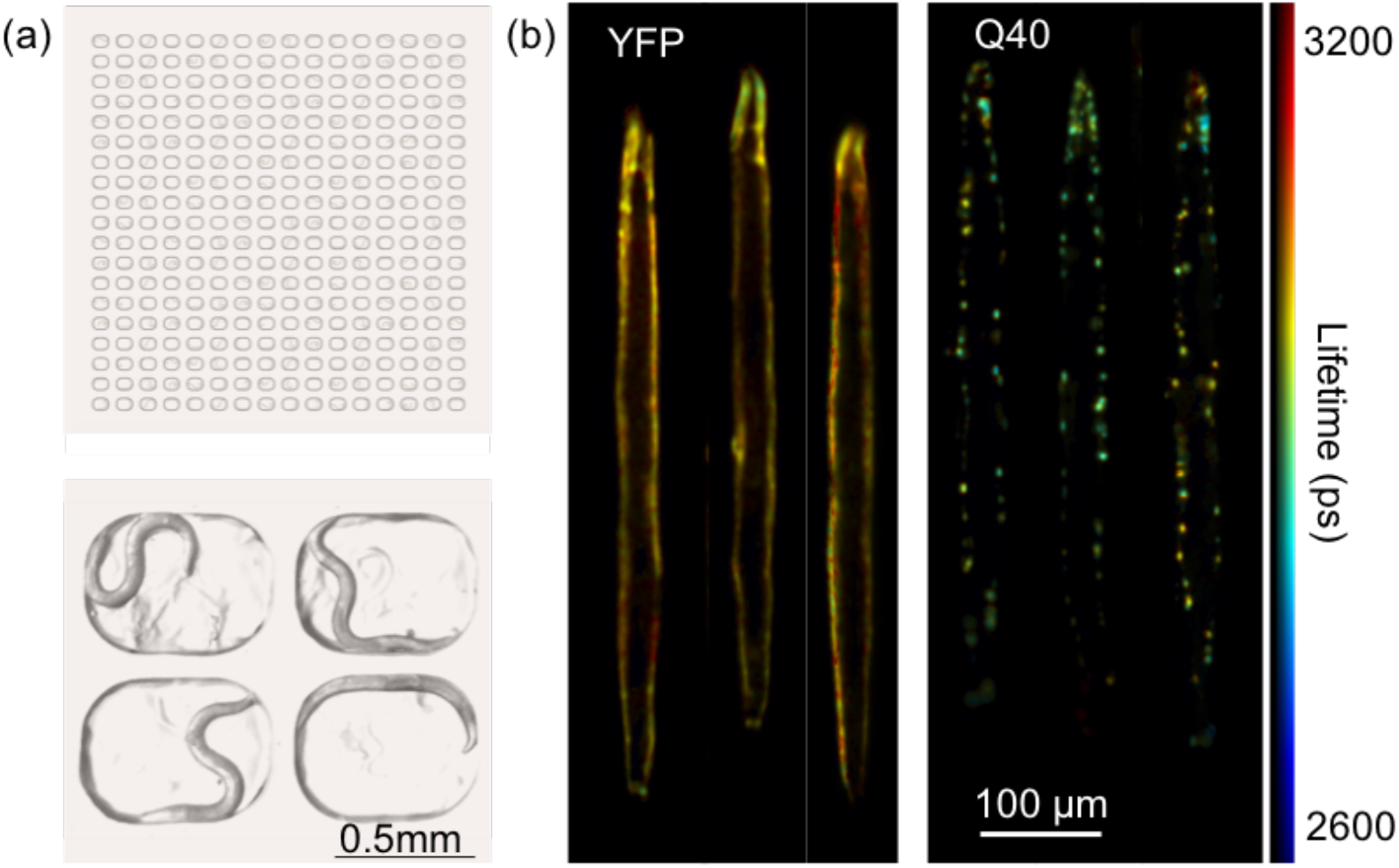
TG-FLIM applied to live *C. elegans* crawling in agarose micro-chambers. **(a)** Schematic of the agarose-based micro-chamber device loaded with *C. elegans* (top) and an image of four chambers, each occupied by a single worm (bottom). **(b)** Registered and digitally stretched fluorescence lifetime maps of live crawling *C. elegans*. Shown here are example images of YFP and Q40 worms at day 3 of adulthood. For each worm, 10 sequential FLIM acquisitions were carried out (equalling a total acquisition time of ~5 s, therefore comparable to that used for anaesthetised worms ~4.2 s), and the lifetime maps were averaged after digital stretching

### Concluding remarks

In conclusion, we have presented here a method for the functional study of protein aggregation in live *C. elegans*. The method reveals differences in the aggregation kinetics and the nature of the inclusions formed during aging in models of Parkinson’s (α-syn) and Huntington’s (Q40) diseases. We observed that α-syn became localised to inclusions prior to the decrease in fluorescence lifetime that is associated with amyloid formation, suggesting that the mechanism of a-syn aggregation involves the persistence of relatively disordered intermediate species prior to the formation of amyloid structure. We furthermore noticed a considerable spread in the fluorescence lifetimes of aged a-syn worms, suggesting that some animals remain largely unaffected by amyloid aggregation even at old age. By contrast, the data showed that Q40 accumulates completely into amyloid inclusions early in adulthood of the worm population. This approach could be extended to image young Q40 nematodes in larval stages at higher resolution, to find out if similar non-amyloid inclusions comprised of e.g. oligomeric_28_ or liquid-like states_37_ are visible as precursors in this system.

For the longitudinal studies described here, we were able to perform the complete set of experiments in less than 2.5 h for each day of measurement (ca. 20 worms for each of the 3 strains studied here). This fast acquisition time provides a high level of reproducibility and has major advantages for high-throughput studies, as it limits the variability in worm age across the experiment while providing sufficient data to define the lifetime changes precisely. This feature constitutes a significant improvement compared to TCSPC where the measurement of ~60 worms would take over ~10 h on a single imaging day. The method also has much higher throughput than FRAP, providing information about the distribution and the aggregation states of all of the inclusions in the imaged region of the animal in one single acquisition. Crucially, TG-FLIM allows us to confirm the amyloid-like nature of the inclusions based on the ability of the FLIM sensor to distinguish between different forms of aggregation_14_, whereas FRAP informs solely on diffusion, which may be similarly restricted for multiple forms of aggregates. Given its speed, we anticipate that the TG-FLIM method will provide new avenues for high-throughput studies of *in vivo* protein aggregation, e.g. to screen for small molecules with the ability to inhibit this process_38_.

We have in addition demonstrated two novel analytical approaches in combination with fast TG-FLIM to image moving nematode worms. The first one uses non-rigid transformation for gate realignment and correction of motional artefacts within the TG-FLIM dataset, which we note would not be possible with TCSPC measurements. The second uses a pseudo-templating method to allow for FLIM map averaging, and for the alignment of all the imaged worms from a given population for easy evaluation of variability and phenotypical properties, e.g. the size of the worms and the spatial distributions of the fluorescent marker.

The results set the stage to apply the fast TG-FLIM approach to more advanced types of chamber devices that could support long-term culturing of *C. elegans*, and enable protein aggregation in the same individual worm to be tracked over time. Agarose micro-chambers have been used to follow the development of *C. elegans* larvae_39_, but microfluidic devices may be necessary for long-term growth, providing a continuous supply of nutrients and separation of offspring as described e.g. by Cornaglia *et al*._40_. The combination of our fast TG-FLIM approach with microfluidic or micro-chamber devices, such as the one that we have presented here, with fully automated data acquisition will greatly improve the throughput of the method compared to manual scanning, as used in the current study. This approach would constitute an invaluable tool for performing very large functional screens for genetic modifiers or compounds that perturb protein aggregation. Optical sectioning capabilities_41_ can also be added to the imaging procedure in order to reveal the 3D organisation of the system, which will provide important additional information e.g. on the subcellular localisation of protein inclusions.

Finally, the fast FLIM imaging demonstrated here can reach a speed of 2 FLIM frames per second (as demonstrated by the 7 gates imaging experiments), and hence can lead to the observation of biologically relevant protein-protein interactions and biosensor dynamics in freely moving worms. Therefore, the use of this fast FLIM method opens up important avenues for time-dependent functional studies using other biosensors, for example to probe Ca^2+^ levels for monitoring neuronal activation while having simultaneous read-outs of the associated behavior of the worms.

## Methods

### TG-FLIM imaging

The TG-FLIM system was set up on an Olympus IX83 frame. The laser source was a super-continuum laser source Fianium SC400-4, spectrally selected by a combination of two linear variable filters and set up for epifluorescence excitation by focussing the beam in the back focal plane of the microscope objective. The fluorescence image was relayed onto the photocathode of a high-rate imager (HRI, Kentech) and its phosphor screen was re-imaged onto the camera sensor (PCO pixelfly, USB, PCO). The magnification of the relays was set up such that the element size of the HRI matched the resolution of the microscope and that of the camera pixel size in 2×2 binning mode. Details of the architecture of the microscope are shown in **Supplementary Figure 1**. Imaging of worm heads was performed using a 40X objective (Olympus UApoN340 40X NA 1.35) and that of entire worms in imaging chambers was performed using a 10X (Olympus PlanFLN 10X NA 0.3) objective. The excitation wavelength was selected to be 516 nm (10 nm bandwidth) and the fluorescence was detected using a 550/49 (Semrock) filter. The instrument response function (IRF) was measured by taking an acquisition of a 1 mM solution of Erythrosin B (Sigma-Aldrich) solution in water. Additionally, the microscope was equipped with a CMOS camera (Blackfly S, BFS-U3-51S5M-C, FLIR) for bright-field measurement.

### FLIM lifetime image reconstruction

FLIM reconstruction was performed using the FLIMfit_42_ package (v4.12.1) from the Open Microscopy Environment (OME). Data analysis was performed by subtracting a background image acquired separately and spatially-varying the IRF reference reconvolution using a lifetime of ~90 ps. A single exponential decay was fitted to the data on a single-pixel basis.

### *C. elegans* culturing and sample preparation

Nematodes were grown under standard conditions on nematode growth media (NGM) plates seeded with *Escherichia coli* OP50 at 20 °C. Worm strains used in these experiments were AM134 expressing YFP, AM141 expressing glutamine_40_-YFP (Q40)_4_, and OW40 expressing α-synuclein-YFP (α-syn)_5_, all under control of the unc-54 promoter to drive expression in body wall muscle cells. Age-synchronized worm populations were generated by a 4 h synchronized egg lay, and animals were transferred to NGM plates containing 75 μM 5-fluoro-2’-deoxyuridine (FUDR, Sigma) at the fourth larval stage to inhibit the generation of offspring.

For imaging, worms were transferred to a drop of M9 buffer (3 g L^−1^ KH_2_PO_4_, 6 g L^−1^ Na_2_HPO_4_, 0.5 gL^−1^ NaCl, 1mM MgSO_4_) containing NaN_3_ as an anaesthetic on a freshly prepared pad of 2.5 % agarose. A cover slip was delicately placed on top and the sample was inverted for imaging on the inverted microscope.

### Imaging chambers

A silicon master with the desired microstructures was fabricated with standard photolithography techniques. The device was designed as an array of 19 x 16 micro-chambers spaced by 350 μm in both directions. The chambers were shaped as rounded rectangles (700 μm x 500 μm x 80 μm depth) to fit within the field of view of the microscope when using the 10X objective (898 μm x 671 μm). Micro-chamber devices were made by pouring 5% high-melting agarose in S-basal buffer (5.85 g L^−1^ NaCl, 1 g L^−1^ K_2_HPO_4_, 6 g L^−1^ KH_2_PO_4_, 5 mg L^−1^ cholesterol) onto the master in a petri dish, and carefully cutting it out after solidification. To allow for sufficient oxygen supply during prolonged imaging times, slits were cut on the back of the device. A drop of *E. coli* OP50 resuspended in Luria Broth medium (10 g L^−1^ bacto-tryptone, 5 g L^−1^ bacto-yeast, 5 g L^−1^ NaCl) was applied onto the device, after which it was left to dry. Worms were washed off NGM plates with M9 buffer (3 g L^−1^ KH_2_PO_4_, 6 g L^−1^ Na_2_HPO_4_, 0.5 g L^−1^ NaCl, 1mM MgSO_4_) and allowed to sediment, after which a ~100 μL drop of solution containing the worms was put onto the device. We observed that the worms tended to swim towards the bottom of the chambers as the drop was drying. However, spreading them with a platinum wire ensured a more homogeneous distribution with most chambers containing a single worm, or being empty. In our hands, the optimum chamber filling was achieved by loading ca. 150-200 worms onto the device, which contains 304 micro-chambers. As soon as the device was dry, a coverslip was put on top and a glass slide at the bottom, after which the sample was imaged in an inverted fashion.

### Correction of motion artefacts and digital stretching

The correction of motion artefacts was performed by using a non-rigid transformation using the MATLAB B-spline image registration written by Dirk-Jan Kroon (MathWorks File Exchange) and initially implemented by Rueckert *et al*._43_. The brightest image of the fluorescence decay was used as a template. Each gate was re-scaled by histogram equalization prior to registration. The transformation obtained for each re-scaled gate was then applied to the corresponding original data. The registered dataset was then saved as OME-tiff for subsequent FLIM analysis.

The digital stretching was performed as followed: each total intensity image was re-scaled by histogram equalization followed by binarization (Otsu thresholding) and active contouring in order to obtain a faithful outline of the worm. The backbone of the worm was obtained by skeletonization and then used to calculate the position of the center of the worm along the geodesic line of the backbone. The backbone was subsequently smoothed by undersampled cubic spline interpolation. The coordinates along the geodesic line were used to estimate the angle of the worm at every point along the backbone and to rotate the image of the worm. For each position the section of the worm image normal to the backbone was obtained and the sections were used to reconstitute the stretched image of the worm. The transformation obtained this way could then be applied to the fluorescence lifetime image and total intensity map.

## Acknowledgements

We would like to thank Dr Sean Warren and Dr Ian Munro for maintaining and updating the FLIMfit package, Dr Christopher Rowlands and Craig Russell for fruitful conversations about the project, and the Caenorhabditis Genetics Center for strains. GSKS and CFK acknowledge funding from the UK Engineering and Physical Sciences Research Council (EPSRC) (grants EP/L015889/1 and EP/H018301/1), the Wellcome Trust (grants 3-3249/Z/16/Z and 089703/Z/09/Z) and the UK Medical Research Council, MRC (grants MR/K015850/1 and MR/K02292X/1), MedImmune (Astra-Zeneca) and Infinitus (China), Ltd. This project has furthermore received funding from the European Union’s Horizon 2020 research and innovation programme under Grant Agreement No. 722380 (to CP), the Netherlands Organisation for Scientific Research (Rubicon fellowship 680-50-1503 to TS), the European Molecular Biology Organisation (long-term fellowship ALTF 72-2015 to TS) and the Centre for Misfolding Diseases of the University of Cambridge (TS, MP, CMD and MV). RFL also acknowledges the support of the UK Biotechnology and Biological Sciences Research Council (BBSRC) (grant BB/P027431/1).

